# Profound lipid dysregulation in mutant TDP-43 mice is ameliorated by the glucocerebrosidase 2 inhibitor ambroxol

**DOI:** 10.1101/2022.08.30.505901

**Authors:** Sophia Luikinga, Alexandre Henriques, Shyuan T. Ngo, Thusi Rapasinghe, Jean-Philippe Loeffler, Michael Spedding, Bradley J. Turner

## Abstract

The importance of dyslipidemia in amyotrophic lateral sclerosis (ALS) patients is increasingly recognised as a potential key mechanism driving disease onset, progression and survival. Evidence in familial ALS models suggests that lipid composition is significantly affected, however clinically relevant models have yet to be investigated. Using a powerful lipidomic approach, we uncover significant dysregulation of glycosphingolipid (GSL) metabolism in both the spinal cord and skeletal muscles of transgenic TDP-43^Q331K^ mice. Treatment with the selective glucocerebrosidase 2 (GBA2) inhibitor ambroxol at symptom onset significantly improved motor and gait functions in TDP-43^Q331K^ mice. Ambroxol treatment preserved motor neurons and neuromuscular junctions which was associated with modulation of GSL metabolism. Our study establishes significant lipid dysregulation in a clinically relevant model of ALS. Importantly, we show positive therapeutic outcomes in a mouse model of TDP-43 proteinopathy, suggesting that ambroxol may be a promising candidate to treat underlying dyslipidemia and symptoms of ALS.

## Introduction

Amyotrophic lateral sclerosis (ALS) is a progressive neurodegenerative disease caused by selective death of motor neurons, resulting in denervation of skeletal muscles and loss of motor control. This eventuates in the inability to swallow, as well as respiratory failure, which causes fatality approximately 5 years after diagnosis. To date, several key genes have been associated with familial ALS, including superoxide dismutase 1 (SOD1), TAR DNA binding protein 43 (TDP-43) and chromosome 9 open reading frame 72 (C9ORF72). However, the majority (~90%) of ALS cases are sporadic and therefore without a known cause, although cytoplasmic TDP-43 is a pathological hallmark. Emerging evidence has linked metabolic deficits to ALS progression and severity [1]. Specifically, abnormalities in lipid metabolism are observed in a high proportion of patients, which seem to contribute to denervation and motor neuron loss [2, 3]. Additionally, hypolipidemia (i.e. low lipid content) is observed in mutant SOD1 mice prior to symptom onset [4].

The brain is one of the most lipid-rich organs and high abundance of lipids is found throughout the central nervous system (CNS). A subclass of lipids that are highly enriched in the CNS are sphingolipids, which are the second most abundant membrane lipid in mammalian cells and are a core structure of cell stability. Besides regulating membrane formation and stability, these sphingolipids are involved in several important cellular processes in the CNS, such as signal transduction and energy homeostasis [5]. Several studies have investigated lipid profiles in ALS patient biofluids and identified dysregulation of sphingolipids, such as glycosphingolipids (GSLs) and sphingomyelin (SM) [6]. Transcriptomic studies have identified glucosylceramide and ceramide as key GSLs that are dysregulated in ALS [7, 8]. An extensive lipidomic study in SOD1^G93A^ transgenic rats has shown little lipid changes in the motor cortex, where drastic changes were found in the lumbar spinal cord lipidome [9]. Furthermore, it has been shown that in mutant SOD1 mice, these GSLs were rearranged prior to symptom onset, indicating that targeting GSL metabolism could be a promising avenue to treat ALS. Indeed, partial prevention of glucosylceramide degradation in SOD1 mice by administration of conduritol B epoxide (CBE) delayed disease onset and improved disease progression in these mice.

CBE is an inhibitor of lysosomal glucocerebrosidase 1 (GBA1), which could be problematic as loss-of-function mutations in GBA1 are causative of Parkinson’s disease [10]. It has been shown that the generic, mucolytic drug ambroxol functions as a selective non-lysosomal GBA inhibitor [11], which was later confirmed to inhibit GBA2 activity without affecting lysosomal GBA1 [12]. This study also shows that chronic treatment with ambroxol in SOD1 mice delayed disease onset and progression. However, SOD1 mutations only represent 10-20% of familial cases and account only for a very small proportion of ALS patients. Therefore, the need of a more clinically relevant model is essential to elucidate lipidomic disturbances, as well as efficacy of ambroxol. Approximately 97% of all ALS cases exhibit TDP-43 pathology, which is considered a hallmark of the disease, while patients and mice harbouring SOD1 mutations lack robust TDP-43 pathology [13]. Recently, transgenic mice expressing mutant TDP-43^Q331K^ were generated and these mice display sporadic features of ALS pathology [14].

Therefore, the aim of this study was to investigate the extent of lipid dysfunction in the lumbar spinal cord, as well as skeletal muscle, of TDP-43^Q331K^ transgenic mice, compared to wildtype controls. We then sought to determine the efficacy of ambroxol on primary motor neurons from transgenic SOD1^G93A^ rats. Lastly, we treated TDP-43^Q331K^ transgenic mice with ambroxol aiming to improve motor function and motor neuron preservation. We demonstrate profound lipid restructuring in both spinal cords and skeletal muscles of TDP-43^Q331K^ mice, highlighting abnormal GSL metabolism. Ambroxol treatment improved survival of motor neurons expressing mutant SOD1 or C9ORF72. Furthermore, we show that chronic ambroxol treatment at symptom onset improved motor function and prevented motor neuron loss and denervation of neuromuscular junctions (NMJs) in TDP-43^Q331K^, consistent with modulation of GSL metabolism. Using multiple and clinically relevant models, we are therefore confident to repurpose ambroxol as a novel potential treatment for ALS.

## Results

### Lipid dysregulation in spinal cords and skeletal muscles of mutant TDP-43 mice

We first interrogated lipid changes in affected lumbar spinal cord and tibialis anterior (TA) muscles of transgenic TDP-43^Q331K^ mice at a symptomatic age (P300), compared to age-matched wild-type (WT) controls, using LCMS targeted lipidomic profiling. Principal component analysis (PCA) of WT and TDP-43^Q331K^ mouse lipidomic profiles showed some separation in spinal cord (Fig 1A). A volcano plot analysis to identify altered lipid profiles performed with fold change (FC) ≥ 1.5 and p-value < 0.05 by t-test revealed 31 significantly changed lipid species in TDP-43^Q331K^ mouse spinal cord, involving 24 enriched and 7 depleted species (Fig. 1B). Heatmap visualization of the top 50 differentially abundant lipid species revealed profound lipid restructuring in spinal cords of TDP-43^Q331K^ mice (Fig. 1C), mainly affecting sphingomyelins (SM), triglycerides (TG) and phosphatidylcholines (PC), which all have several subclasses significantly affected. Raw peak analysis revealed accumulation of all ceramide (Cer) subtypes in TDP-43^Q331K^ mice, where acylcarnitine (Acar) only had one subtype more abundant, dihydroceramide (DhCer) two and GM3 one (Fig 1D). Together, these data strongly suggest impaired ceramide metabolism. Interestingly, Acar levels which link to ceramide metabolism and mitochondrial function, were higher in TDP-43^Q331K^ mice. Thus, ceramide and related GSL metabolism are profoundly dysregulated in the CNS of mutant TDP-43 mice.

**Figure 1.**
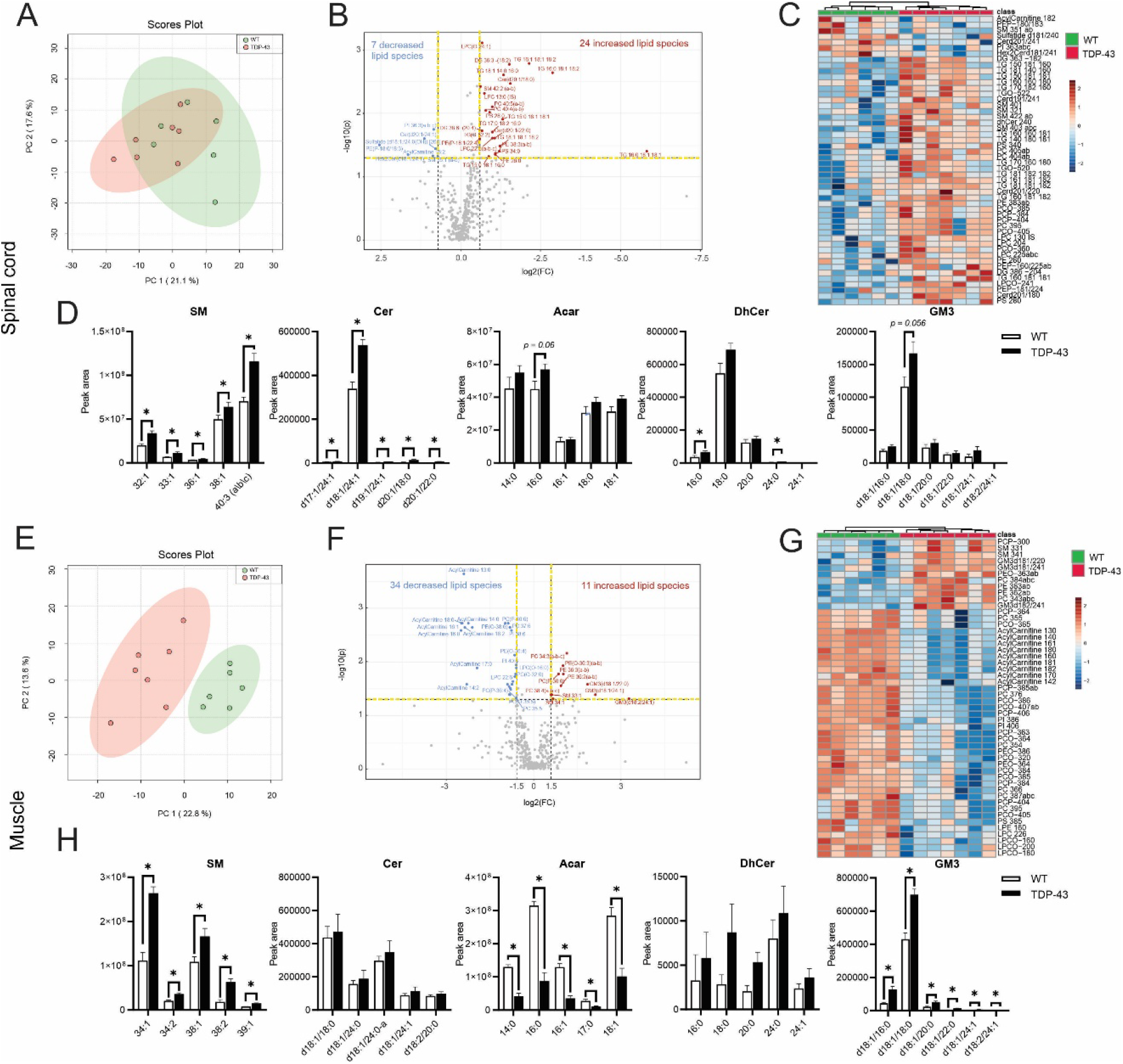
Lipidomic profiles of TDP-43^Q331K^ mice. Principal component analysis of wildtype (WT) and TDP-43^Q331K^ mouse (A) lumbar spinal cord and (E) tibialis anterior (TA) muscle, the explained variances in brackets on the relevant axes. Lumbar spinal cord lipids show some clustering, with relatively large variability, where TA lipidome is more separated. Volcano plot with fold change threshold (X-axis) of 1.5 and *p*-value cut-off at 0.05 (Y-axis) coloured circles represent lipids above these thresholds, in the spinal cord there were 24 decreased and 7 increased lipids in WT vs. TDP-43 mice (B), where in the Tibialis Anterior muscle 34 were decreased and 11 increased (F). Heatmap of targeted LCMS in lumbar spinal cord (C) and TA (G) shows clear separation based on genotype suggesting distinct lipid dysregulation in TDP-43 mouse tissues. Raw peak data visualisation of shows 5 main GSLs thought to be involved in ALS pathogenesis in the spinal cord (D) and the TA (H). All 5 are more abundant in the lumbar spinal cord, with a clear difference between the lipid subtypes, where in the TA Acar is depleted. and tibialis anterior (TA) muscle (B).

In TA muscle, there was a clear dissociation of lipid signatures in TDP-43^Q331K^ mice, compared to WT mice (Fig. 1E). Volcano plot and t-test analysis of TA muscle revealed 45 significantly altered lipid species in TDP-43^Q331K^ mice (Fig. 1F). Notably, Acar and GM3 were clearly dysregulated. Heatmap depiction of the top 50 lipid species dysregulated in TDP-43^Q331K^ mouse TA (Fig. 1G), revealed marked lipid restructuring, reflected by increased abundance of SM, phospholipids (PC, PE) and GM3, and decreased abundance of Acar. A strikingly different lipid profile to spinal cord was observed in TA muscle, with accumulation of SM and depletion of Acar, again highlighting ceramide metabolism (Fig. 1H), while all subtypes of Acar levels were severely depleted in muscles of TDP-43^Q331K^ mice. GM3 abundance was again increased in muscles of TDP-43^Q331K^ mice. Interestingly, ceramide accumulation was not evident in skeletal muscle of mutant TDP-43 mice. Further observation of these lipidomic results suggested that lipid composition of skeletal muscle in TDP-43 mice was more affected than that of spinal cord, evidenced by the clearer separation of groups in the PCA plots (Fig 1A & E).

Untargeted lipidomic analysis was also performed for both lumbar spinal cord and TA muscle with sphingolipids mostly affected in TDP-43^Q331K^ mice. A total of 80 lipids were dysregulated in lumbar spinal cord, with 210 lipids altered in skeletal muscle (Supplementary Fig 2), again indicating that skeletal muscle is more affected by lipid dyshomeostasis in mutant TDP-43 mice.

We uncovered a lipidomic signature in spinal cord and skeletal muscles of TDP-43^Q331K^ mice showing major dysregulation in the GSL pathway, with ceramide central. Therefore, we aimed to seek out the importance of lipid metabolism on motor neuron survival, function and TDP-43 mislocalisation in ALS. We targeted sphingolipid metabolism by inhibiting glucocerebrosidase 2 (GBA2), an enzyme responsible for synthesising GlcCer into ceramide, using ambroxol, a selective GBA2 inhibitor. A perixosome proliferator-activated receptor γ (PPARγ) antagonist T0070917, was used to investigate the molecular mechanism of ambroxol in lipid metabolism. PPARγ regulates fatty acid storage and glucose metabolism and is a key receptor for mitochondrial function. Further, GM1 and oligo-GM1 were used to investigate the neuroprotective effects of these compounds as these seem to be central in sphingolipid metabolism.

### Ambroxol improves motor neuron survival, neurite network and prevents TDP-43 mislocalisation *in vitro*

The effects of ambroxol on primary motor neurons challenged with glutamate, an ALS relevant stressor, were determined. One way ANOVA, followed up with *post hoc* analysis revealed that 10 μM of ambroxol was sufficient to improve survival (*p*<0.05) of primary motor neurons prepared from WT rats, compared to glutamate alone (Fig 2A & B). Riluzole alone also improved survival (p<0.05), as well as combined with ambroxol (*p*<0.05). Next, we examined the impact of ambroxol on the integrity of neurites in motor neurons. Ambroxol, riluzole or both drugs in combination rescued neurite network disruption following glutamate challenge (*p*<0.05) (Fig 2B). Lastly, we assessed the effects of ambroxol on cytoplasmic TDP-43 accumulation, a pathological hallmark of ALS. Ambroxol, riluzole or both drugs together prevented TDP-43 mislocalisation induced by glutamate exposure in primary motor neurons (*p*<0.05) (Fig 2B). Together, we show here that ambroxol improves function of injured motor neurons.

**Figure 2.**
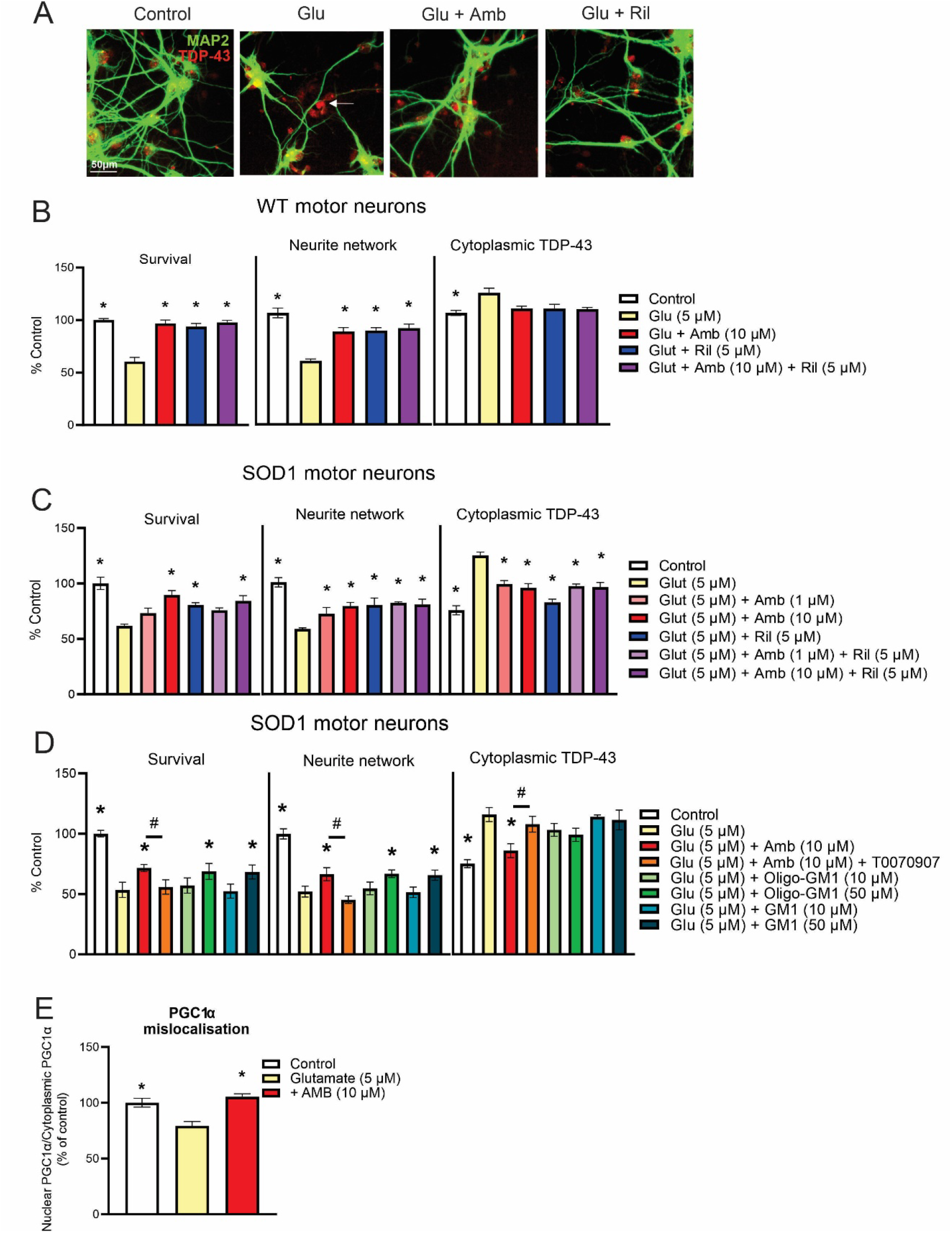
*In vitro* effects of lipid metabolism and efficacy of ambroxol. Representative image of cultured rat primary motor neurons with clear TDP mislocalisation after glutamate challenge as indicated by the white arrow. Neurite networks stained with MAP-2 in green are also clearly affected. Effects of ambroxol (10 μM) with and without riluzole (5 μM) on primary motor neuron cultures challenged with glutamate (20 μM) for 20 min. Glutamate alone caused motor neuron death, disrupted neurite network and TDP-43 mislocalisation. Ambroxol prevented motor neuron loss, preserved neurite network (B). In transgenic SOD1^G93A^ rat primary motor neurons, ambroxol exerted similar neuroprotective effects (C). Analysis of other important lipid components on transgenic SOD1^G93A^ rat primary motor neurons indicates that when ambroxol is given with a PPARγ antagonist, survival and TDP-43 mislocalisation effects are diminished. Treatment with higher concentrations of GM1 (50 μM) shows improved survival, neurite network and prevents TDP-43 mislocalisation (D). Mislocalisation of PGC1α in transgenic SOD1^G93A^ rat primary motor neurons was also rescued by ambroxol, indicating mitochondrial importance in the working mechanism of ambroxol (E).

Next, we established the effects of ambroxol on primary motor neurons cultured from transgenic SOD1^G93A^ rats. We showed clear glutamate-mediated excitoxicity induced in these motor neurons which was antagonised by ambroxol (1 and 10 μM) and riluzole (5 μM) using three endpoints (survival, neurite network, cytoplasmic TDP-43, p<0.05, Fig 2C). The effects of ambroxol were observed at similar concentrations that were reported to increase NMJs in spinal explants/myoblasts [15]. We then investigated the effects of lipid importance in these glutamate challenged SOD1^G93A^ motor neurons. The effects of ambroxol were abolished by the PPARγ antagonist T0070907 (10 μM), while GM1 (50 μM) was protective against neuronal cell death and neurite depletion (Fig 2D). In additional experiments, ambroxol (10 μM) protected against the effects of glutamate on PGC1∝ mislocalisation (Fig 2E). Thus, ambroxol is neuroprotective in models of familial ALS and excitoxicity implicated in sporadic ALS, and these data support the importance of lipid metabolism in mediating these beneficial effects.

### Ambroxol treatment improves motor function in TDP-43^Q331K^ mice

Since we observed positive treatment effects *in vitro*, we were confident to use ambroxol *in vivo*. We administered 3 mM ambroxol in drinking water to TDP-43^Q331K^ mice from disease onset (P60) and examined motor activity. Analysis of motor behaviour using a rotarod showed a deficit in TDP-43^Q331K^ mice, compared to WT controls starting at P60 (Fig. 3A). TDP-43^Q331K^ mice developed a slow and gradual motor deficit over 8 months. Using 3-way RM ANOVA, we found a main effect of latency to fall over time F_(32,1120)_=26.13, *p*<0.001. There was a main effect of genotype F_(1,35)_ = 19.3, *p*<0.01, where TDP-43^Q331K^ mice significantly performed worse. Importantly, there was also a main effect of ambroxol treatment F_(1,35)_=3.8, *p*<0.05, demonstrating that ambroxol improved motor function of TDP-43^Q331K^ mice, compared to vehicle TDP-43^Q331K^ mice. Ambroxol efficacy was also confirmed by locomotor cell analysis (Fig. 3B). While voluntary movement was normal at P60, independent t-test indicated that TDP-43^Q331K^ mice showed a significant deficit at P270 (p<0.05), compared to WT controls. TDP-43^Q331K^ mice treated with ambroxol showed improved voluntary movement, compared to vehicle TDP-43^Q331K^ mice (p<0.05). Thus, ambroxol administration improves the phenotype of TDP-43^Q331K^ mice across different locomotor tests.

**Figure 3.**
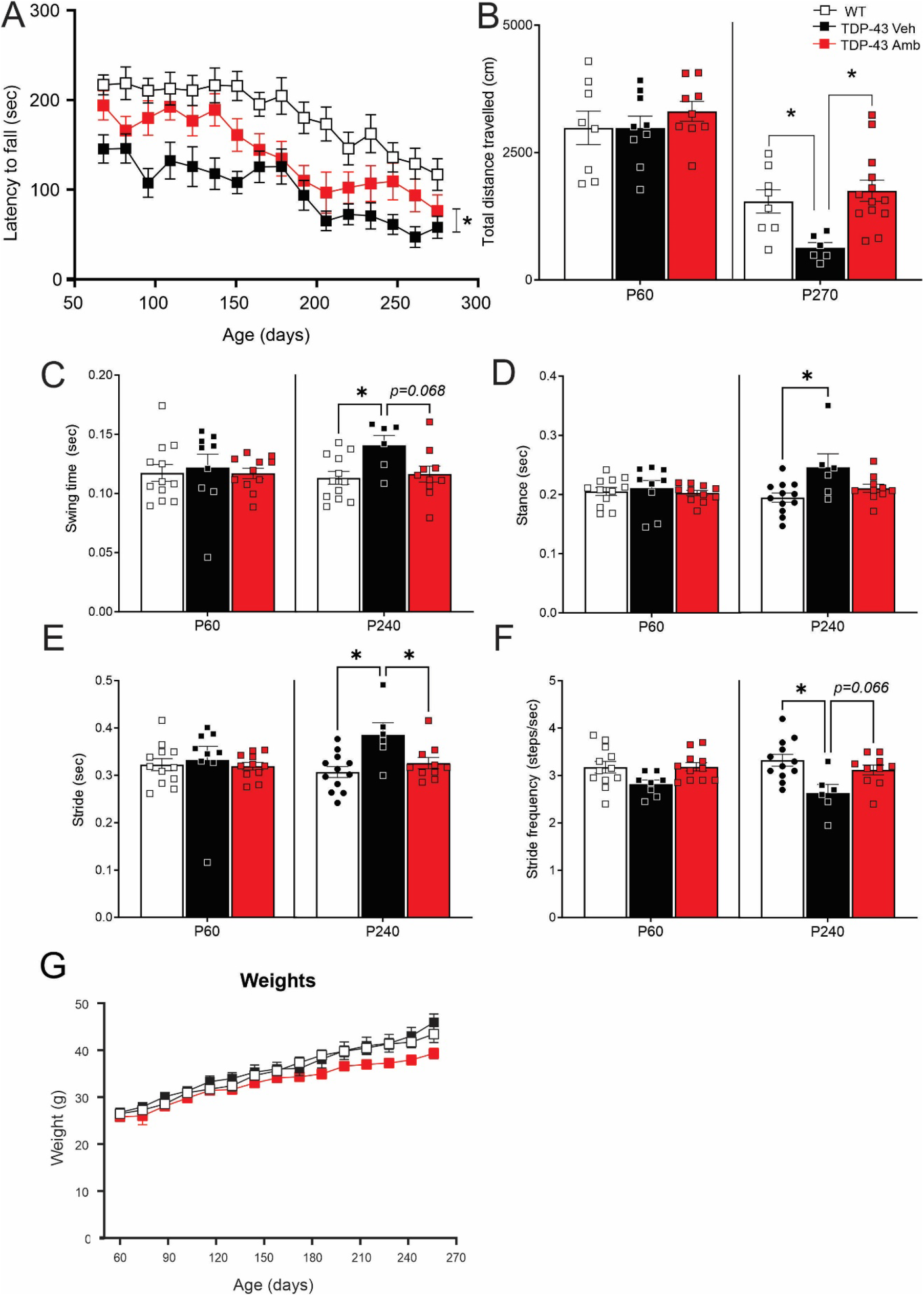
Ambroxol treatment improves the motor phenotype of TDP-43^Q331K^ mice. (A) Rotarod analysis of wildtype (WT) and TDP-43^Q331K^ mice treated with and without ambroxol. Ambroxol significantly improves rotarod performance in TDP-43^Q331K^ mice, compared to vehicle TDP-43^Q331K^ group. (B) Locomotor cell analysis of mice shows a deficit in spontaneous locomotion in vehicle TDP-43^Q331K^ mice which is significantly rescued by ambroxol at P270. Digi-gait analysis of mice reveals abnormal (C) swing time, (D) stance, (E) stride and (F) stride frequency in vehicle TDP-43^Q331K^ mice which is ameliorated by ambroxol treatment at P270. (G) Body weights of mice over time. *p<0.05, 3-way RM ANOVA or 1-way ANOVA, *n*=6-12 mice/group.

We next assessed gait function in mice using DigiGait™ analysis. A full report on DigiGait™ analysis can be found in Supplementary Table 2. At P60, stride, stride frequency, stance and swing times were similar in TDP-43^Q331K^ and WT mice (Fig. 3C-F). At P240 however, TDP-43^Q331K^ mice revealed deficits in all these gait measures (*p*<0.05, relative to WT controls). Ambroxol administration significantly improved stride (*p*<0.05) and there was a positive trend observed between TDP-43 vehicle and ambroxol treated mice in stride frequency and swing time. It is important to note that these gait measures in ambroxol treated TDP-43 ^Q331K^ mice do not statistically differ from WT mice, indicating that ambroxol prevents gait deterioration in TDP-43 mice. Lastly, RM ANOVA did not show a significant effect of ambroxol on body weight of mice (*p>*0.1, F>1 Fig 3G).

### Ambroxol preserves motor neurons and NMJs by upregulating gangliosides in TDP-43^Q331K^ mice

To determine the neuroprotective effects of ambroxol, we quantified motor neuron degeneration in lumbar spinal cords of TDP-43^Q331K^ mice. Mutant TDP-43 mice showed a ~10% loss of spinal motor neurons, compared to WT mice (*p*<0.05) (Fig. 4A,B). Ambroxol treatment significantly prevented motor neuron loss in TDP-43^Q331K^ mice, compared to vehicle controls (p<0.05). Next, we assessed the effects of ambroxol on muscle denervation. Analysis of acetylcholine receptor 1α (*AChR1α*) levels in TA muscles (known to increase with denervation [16]) revealed upregulation in TDP-43^Q331K^ mice (p<0.05) by qPCR, which was ameliorated by ambroxol (Fig 4C). Gene expression of other denervation markers *MUSK* and *Atrogin* were unchanged in TDP-43^Q331K^ muscle (Supplementary Figure 3D). NMJ integrity was next assessed in TA muscles. TDP-43^Q331K^ mice showed a significant ~50% loss of innervated NMJs revealed by co-staining of presynaptic markers and α-bungarotoxin (Fig 4D,E). Denervation was significantly prevented in TDP-43^Q331K^ mice treated with ambroxol, compared to vehicle TDP-43^Q331K^ mice (p<0.05).

**Figure 4.**
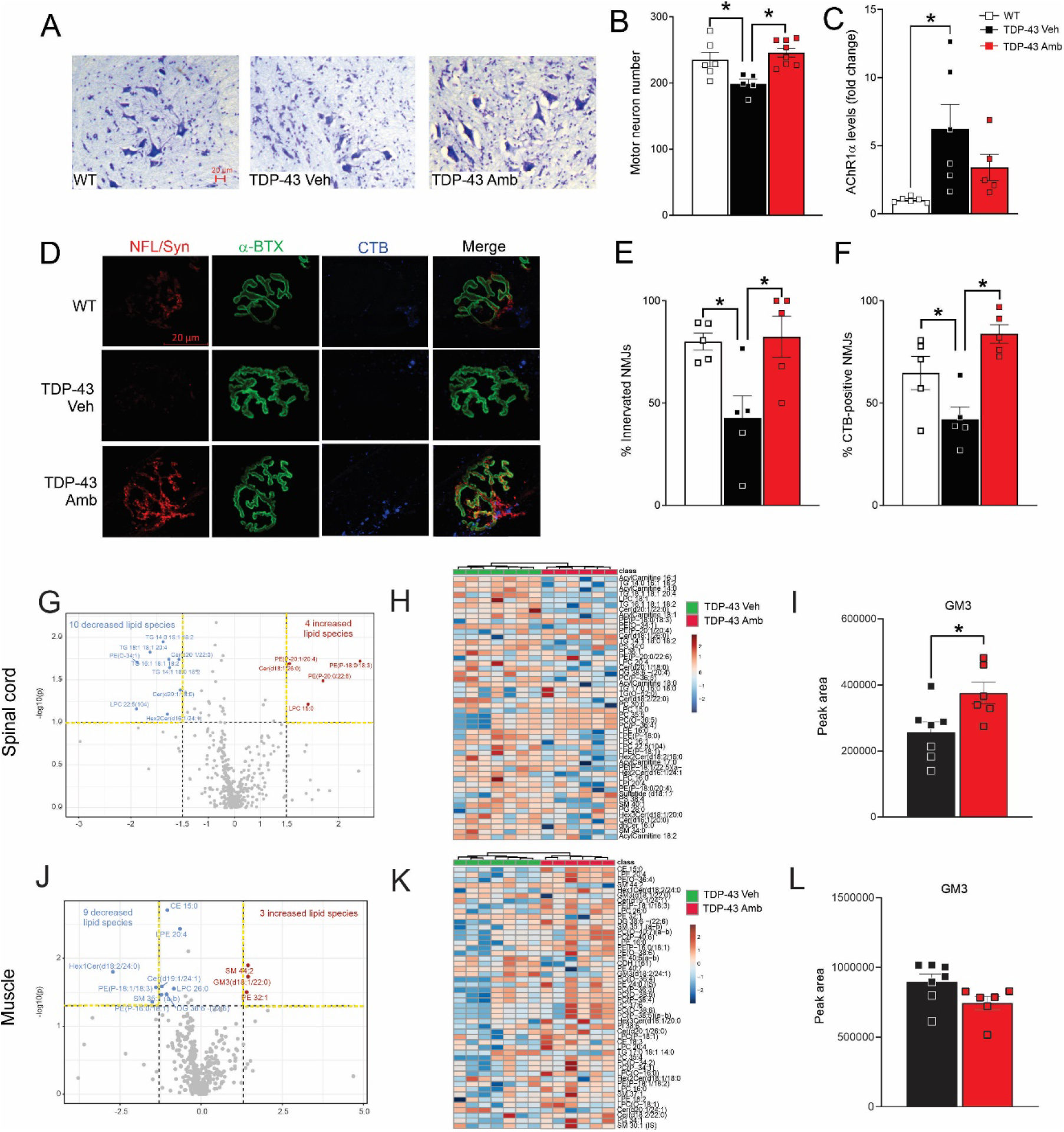
Effects of ambroxol on motor neuron pathology, glycosphingolipids and lipidomic profiles in TDP-43^Q331K^ mice. (A) Representative images of ventral horns of lumbar spinal cords stained with cresyl violet with (B) motor neuron counts showing neuroprotective effects of ambroxol. (C) qPCR analysis of AChR1α expression levels in tibilias anterior (TA) muscle. (D) Representative images of neuromuscular junctions of TA muscle stained for neurofilament/synaptophysin (red), α-BTX (green) and CTB gangliosides (blue). Quantitative analysis of (E) NMJ innervation and (F) CTB abundance shows ambroxol preserves peripheral synapses and gangliosides. Plot generated by Metaboanalyst of one-way ANOVA of untargeted lipidomics (G & H). In the spinal cord 14 significant features were identified and in the TA 12. Raw peak analysis of GM3 in Spinal cord (I) and TA (L) shows higher GM3 abundance in ambroxol treated TDP-43 mouse spinal cord.. *Denotes post hoc significant effect *p* < 0.05, *n*=5-7 mice/group.

To confirm that ambroxol enhances GSL synthesis *in vivo*, we employed immunohistochemical and lipidomic approaches. Distribution of gangliosides (GM1a and others) at NMJs was studied using cholera toxin subunit B (CTB) which binds to gangliosides in neuronal membranes and has the highest affinity to GM1 [17]. CTB labelling was detected at the presynaptic side of NMJs in WT mice which was disrupted in TDP-43^Q331K^ mice (p<0.05) (Fig. 4D,F), consistent with depletion of gangliosides (at least GM3) in mutant TDP-43 mice. Ambroxol treatment significantly restored CTB staining and thus ganglioside levels (mainly GM1) at NMJs, correlating with preserved neuromuscular innervation and function. Next, we profiled lipid changes in spinal cord and TA muscles of TDP-43^Q331K^ mice treated with ambroxol using LCMS targeted lipidomics. In spinal cord, there was a small effect of ambroxol on lipid composition, with a large overlap in the PCA plot (Supplementary Fig 4F) and a total of 14 lipids significantly dysregulated by volcano plot (Fig 4G). However, heatmap analysis shows a clear clustering based on ambroxol treatment. Abundance of several key lipids species, such as Acar, Cer and PE were altered by ambroxol, indicating that ambroxol modulates lipid dyshomeostasis in spinal cords of TDP-43^Q331K^ mice (Fig 4H). Similar to the spinal cord, the effects of ambroxol on lipid composition in TA muscles was small, but observable. A total of 11 lipid species were found to be changed by ambroxol shown by volcano plot (Fig 4J). Heatmap analysis showed a clear clustering based on ambroxol treatment. Key lipids affected by ambroxol in muscle were SM and GM3, as observed by volcano plot and heatmap analysis (Fig 4J,K). Importantly, ambroxol restored at least one subtype of SM and GM3 (*p*<0.05, Fig 4L). When analysing the impact of ambroxol using untargeted lipidomics, the effects of ambroxol on lipid composition was more profound. Ambroxol significantly restructured lipid composition in both spinal cord and muscles of TDP-43^Q331K^ mice. That is, a total of 30 lipids were significantly altered by ambroxol in spinal cord indicated by volcano plot (Supplementary Fig 4B). On the other hand, 39 lipids were significantly changed by ambroxol in TA muscles, with the majority increased (Supplementary Fig 4E). Together, this suggests that ambroxol has the ability to restructure lipids in the CNS and skeletal muscle, indicating that motor improvements, as well as prevention of muscle denervation by ambroxol was likely driven by modulating lipid metabolism.

Some lipids that were found to be dysregulated in TDP-43^Q331K^ mice, such as PE DG and ceramide, are also known to play an important role in autophagy [18], a cellular function known to be disrupted in ALS. Therefore, we analysed key autophagic marker proteins (LC3-I/II, p62 and VDAC1) in spinal cords, however these were not modulated by ambroxol treatment in TDP-43^Q331K^ mice (Supplementary Fig 3A & B). We also assessed gene expression of *p62*, *CathD, CathZ, GNPTAB, Beclin1* and *ULK1*, but did not detect any changes induced by ambroxol (Supplementary Fig 3C).

Lastly, the profound effects of ambroxol on SM (Fig 1) raised a question about myelin integrity in the spinal cord. Therefore, we employed SCoRE on spinal cord sections. Analysis of intensity of myelin in the corticospinal tract (CST) using SCORE imaging revealed no differences in myelin density at P60 or P210 (*p*s >0.05, F<1) (Supplementary Fig 3E & F).

Together, ambroxol treatment protects both motor neuron cell bodies and peripheral synapses, consistent with improved motor function.

## Discussion

These results confirm and extend mounting evidence using metabolomic studies [7, 8, 19, 20] that sphingolipid metabolism is profoundly disrupted at a very early disease stage in animal models of ALS; these changes also occur in patients [7, 8, 20]. Many experimental therapeutic approaches have been tried in ALS, with almost negligible success, so the identification of novel therapeutic targets linked to key aspects of the disease state are of crucial importance. Here, we show that mutant TDP-43 mice have profoundly dysregulated lipid and sphingolipid metabolism. GSL metabolism has been previously been shown be critically modified in mutant SOD1 mouse models and inhibition of glucosylceramide synthase (GCS, also known as UGCG) was shown to be profoundly deleterious for denervation and NMJ function [7, 20], while inhibition of GCase [19] was beneficial. Furthermore, GBA2 activity is profoundly (8-10-fold) increased in spinal cord at the very early stages of disease in mutant SOD1 mice, which would break down complex membrane GSLs, such as GM3 (sialyl-lactosylceramide) and GM1 (monosialotetrahexosylceramide) to equivalent ceramides [15]. In this study, and that of Bouscary et al, (2019), GM1 was shown to be a critical component of NMJs, lost at the very first stage of denervation, presumably caused by the increase in GBA2 activity. GM1 represents >10% of human brain gangliosides, and is a powerful neurotrophin, being required for a NGF-TrkA-GM1 complex (reviewed by [21]), but also acting by the BDNF neurotrophic cascade [22]. GM1 is a cellular target for multiple enveloped viruses, and also for cholera toxin, which is used to label GM1 [23], although the selectivity is not absolute. The sialic acid confers structural rigidity on the trisaccharide chain, crucial for cellular recognition. The sialic acid of GM1, target for viruses, was mutated in hominids about 2.5 million years ago, perhaps because of a virus, in that humans lost CMP-Neu5Ac hydroxylase (CMAH) gene which changes running endurance, and neuromuscular metabolism, at a time when humans evolved to run, with associated metabolic changes, probably associated with BDNF [24] [25]. Purified GM1 has been used to treat peripheral neuropathies [26]. In our lipidomic analysis, we were unable to detect GM1, most likely since the abundance in whole tissue is low, compared to other lipids. We are however, confident that our CTB labelling results are robust and we can conclude GM1 along the motor axons in the NMJ is depleted. As a consequence of the dysregulated lipids observed, it is highly likely that ceramide could accumulate as a direct result. Increased ceramide accumulation is a critical intermediate for multiple metabolomic changes, modifying mTORC1, increasing apoptosis in multiple cell types, and also downregulating glucose transporters [27]: forcing additional lipid metabolism, affecting acyl carnitine transport. The question therefore arises whether the increases in GCS (UGCG) activity occurring in ALS patients and mouse models [7, 8] and GBA2 activity induction in animal models [12] are compensatory or deleterious. We consider that the increase in GCS activity may be compensatory to restore levels of GM2 and GM1a, whereas the increase in GBA2 activity may be deleterious. Additionally, here we show that the abundance of SM in the spinal cord seems to be greater in the TDP-43 mouse. Interestingly, increased SM in the CNS has been shown to be associated with heightened intracellular calcium levels [28]. Elevated intracellular calcium levels could cause neurons to be hyperexcitable, a phenomenon that is detected in the cortex prior to lower motor neuron degeneration [29]. However, based on the current data, we are unable to determine if this SM accumulation precedes hyperexcitability.

Since we found these major sphingolipid disruptions in tissues of TDP-43 mice, we aimed to further investigate the importance of these systems *in vitro* and sought to determine if we can manipulate sphingolipid metabolism using the GBA2 inhibitor ambroxol. We examined the neuroprotective role of ambroxol and GM1, and its oligosaccharide (to eliminate interference from any surfactant effects from the GM1 lipid side-chain), in primary motor neurons subjected to an excitotoxic insult with glutamate. In these experiments, 10 μM ambroxol promoted motor neuron survival and neurite network stability in the presence of glutamate; riluzole was also protective, and the effects of riluzole were not blocked by ambroxol, which is important for clinical trial considerations. Both drugs blocked glutamate-induced TDP-43 pathology, which is relevant to the effects in TDP-43^Q331K^ mice. These effects were presumably due to GBA2 inhibition. In motor neurons from SOD1^G93A^ mice, similar effects were observed, except that ambroxol was active from 1 μM, while still preventing glutamate-induced TDP-43 pathology. The protective effects of ambroxol were entirely blocked by the PPARγ antagonist, T0070917, indicating a potential effect on mitochondrial metabolism, and furthermore, ambroxol improved nuclear translocation of PGC1α in motor neurons, further indicating an association between ambroxol and pro-mitochondrial pathways (PPARγ/PGC1α). Pioglitazone, a PPARγ agonist, has been shown to increase survival in SOD1^G93A^ mice [30], and PPARγ has been proposed to increase oxidative phosphorylation, and mitochondrial biogenesis, while increasing antioxidant defense [31]. GM1 and oligo-GM1 at 50μM, but not 10μM, protected SOD1^G93A^ motor neurons from glutamate-induced toxicity: these concentrations were previously neuroprotective in other systems [32]. Since proven as a potential therapeutic *in vitro*, we were confident to treat the TDP-43^Q331K^ mouse that best represents sporadic ALS preclinically [14].

We show that ambroxol slowed disease progression in this mouse model. While this mouse only has a mild ALS-like phenotype, chronic oral ambroxol treatment improves motor function as measured by rotarod, locomotor cell and Digi-gait analysis. Furthermore, ambroxol exerted major metabolic effects in this clinically relevant model of ALS: with significant increases in GM1 (as measured at the NMJ) and GM3 (d18:2/24:1) in spinal cord, but not muscle, associated with ameliorations in function; these effects are compatible with inhibition of GBA2. There was also a reduction of some acylcarnitines in spinal cord, and an increase on phoshatidylinositol (PI, 36:4 and 38:5), phosphatidyl serine (PS, 36:1; 36:2), lysophosphatidyl choline (LPC, 16:0, 18:0, 18:1), lysophosphatidyl ethanolamine (LPE, 18:0, 18:1). In muscle, there was an increase in multiple species of phosphatidyl ethanolamine (PE). These phospholipids have multiple roles, but are crucially important for mitochondrial function, and PE species have been shown to be essential for mitochondrial biogenesis following exercise training, which disproportionally increases mitochondrial PE compared to other species [33]. Furthermore, deletion of PS decarboxylase, to inhibit synthesis of PE, inhibited the training effects, and eventually caused respiratory failure because of diaphragm muscle failure, implying that an increase in PE will echo beneficial effects on muscle mitochondrial metabolism.

Thus, the effects of ambroxol are widespread, and modify critical pathways for metabolism, which are awry in ALS. These effects may also allow the definition of new biomarkers in humans, which will be explored in clinical trials. In conclusion, GSLs appear to be critical factors in ALS, rationalising therapeutic targeting of GSL metabolism.

## Materials and Methods

### Animals and drug treatment

All animal experiments conformed to the Australian National Health and Medical Research Council published Code of Practice and were approved by the Florey Institute of Neuroscience and Mental Health Animal Ethics Committee (permit number 17-093). Transgenic human TDP-43^Q331K^ mice (B6N.Cg-Tg(Prnp-TARDBP*Q331K)103Dwc/J, line 103, stock number 017933) were purchased from the Jackson Laboratory (Bar Harbour, ME) and maintained as heterozygotes on a C57BL/6NJ background. Non-transgenic C57BL/6NJ littermate mice provided wild-type (WT) controls. Mice were group housed (no more than 6 littermates) in individually ventilated cages (IVC) under standard 12 h light-dark conditions (lights on 0700-1900) at 22°C with regular chow and water available *ad libitum*. Mice were randomly assigned to either ambroxol or vehicle groups. Single housing of animals was avoided, if possible, but sometimes required due to aggressive behaviour of male mice.

Ambroxol hydrochloride (TCI Chemicals, 98% purity) was dissolved and administered *ab libitum* in drinking water at a concentration of 3 mM as previously described [15]. Ambroxol solution was replaced 3 times a week. Vehicle mice received normal drinking water. Treatment started at symptomatic age postnatal day 60 (P60) until study endpoint (P300).

## Behavioural experiments

### Rotarod latency

Locomotor function was assessed weekly using an accelerating mouse Rota-Rod 47600 (Ugo Basile, Italy) as previously described [34]. Prior to testing, mice were familiarised on the rotarod and trained for 2 days with 1 steady session (16 RPM) and two accelerated (4-40 RPM) sessions over 5 min each with a 10 min resting period. During the testing phase, mice were subjected to two 5-minute accelerated sessions (4-40 RPM). The average time latency to fall off the rod was recorded (sec).

### Open field locomotion

Voluntary locomotion was tested monthly using locomotor chambers with infrared detection beams (ENV-510-A, Med Associates, Fairfax, USA). Animals were placed in the middle of the arena and allowed to freely move around. Locomotion was recorded for 30 min and total distance travelled (cm) was measured per animal.

### Gait analysis

Changes in gait were assessed using the DigiGait™ Imaging System (Mouse Specifics Inc., Boston, USA) using methods as described in [35]. Briefly, paws of mice were temporarily stained using a red food dye to enhance detection sensitivity. Mice were placed in the apparatus and allowed to walk at 15 cm/s which was the tolerated speed at later stages of disease. Gait was recorded for a minimum of 10 steps. The videos were automatically digitized, and these images were used to define the area of each paw. Compilation of these digital images generated a set of wave forms per limb. Built in algorithms within the software (Mouse Specifics Inc., Boston, USA) identified 42 different parameters of gait dynamics per limb. Gait parameters altered in SOD1 transgenic mice, identified by [36] were analysed.

### Lipidomics

For lipid analysis mice were killed at P210. Tissue was collected fresh and weighed prior to lipid extraction.

Targeted lipidomic analysis was carried out using an HPLC (Agilent LC 1290) coupled to an Agilent 6490 Triple Quadrupole mass spectrometer (Agilent Technologies, Santa Clara, USA). For HPLC lipid separation, Agilent Poreshell C18 column (100 mm × 1.2 mm, 2.7 μm was used with a binary gradient of solvent A (20:30:50 (v/v/v) Iospropanol:Aceronitrile:water with 10 mM ammonium formate and solvent B (90:0.9:0.1 (v/v/v) Iospropanol:Aceronitrile:water with 10 mM ammonium formate over 15 minute gradient. For each lipid sample, 5 μL aliquots were injected. The column temperature was set at 35°C, and the flow rate of the mobile phase was at 0.4 mL/min. The gradient used was; first a 0-2.7 min gradient elution from 10% B to 45% B which was then increase to 53% B from 2.7 to 2.8 min, then to 65% B from 2.8 to 9 min followed by an increase to 89% B from 9 to 9.1 min. Next, it was increased to 90% B from 9.1 to 10 min followed by an increase to 92% B from 10 to 12 min and an increase to 100% B from 12.1 to 13.9 min. Then, from 13.9 to 15 min B was decreased to 10% for column re-equilibration

Multiple reaction monitoring (MRM) of the Triple Quadrupole MS/MS was used to identify and quantify glycerolipids from each class. The MS was set at positive mode. About 20 to 50 compounds were simultaneously measured every 20 ms dwell time. Each chromatographic peak was about 30 to 45 s wide with a minimum of 12 to 16 data points collected. The MS parameters were optimized with the capillary voltage at 4000 V, fragmentor at 140-380 V, collision voltages at 15-60 V, and collision gas (nitrogen) at 7 L/min. Agilent MassHunter quantitative software (version 6; Mulgrave, Australia) was used to process the acquired data

### Gene expression analysis

Total RNA was extracted from fresh frozen lumbar spinal cord and tibialis anterior muscle using the Qiagen RNeasy kit (Qiagen) according to the manufacturer’s instructions. A total of 500ng cDNA was synthesised from total RNA using the iScript cDNA synthesis kit (Biorad) according to the manufacturer’s instructions. For the qPCR reaction, 5μM of primer and 14 ng of cDNA and 5 μl of 2x PCR Master mix (applied biosystems) was added to each well. Total reaction volume was 10μl. Target genes and primer sequences can be found in supplementary table 1. Relative mRNA expression was normalised to βActin, using wildtype vehicle as control and fold change was calculated using the ΔΔCT method [37].

### Immunohistochemistry

Mice were transcardially perfused with 100 mM PBS (pH 7.4) followed by 4% PFA in 0.1 M phosphate buffer at pH 7.4. Dissected lumbar spinal cords were post-fixed in 4% PFA for 2 h and cryoprotected in 30% sucrose in PBS overnight at 4°C or ontil tissue has sunk. Tissues were embedded in Tissue-Tek OCT compound (Sakura finetek, USA) by freezing on dry-ice. Horizontal 20 μm sections were cut on a semi-automatic cryostat (Leica). For motor neuron counts, sections were stained with 0.5% cresyl violet using a standard protocol. Nissl-positive motor neurons in every third section were counted from a total of 30 ventral horns per mouse (n = 5 mice per group). Motor neurons were identified by neuronal morphology and size exceeding 20 μm diameter with a distinct nucleolar profile, as well as lateral localisation in the ventral horns.

For neuromuscular junction (NMJ) analysis, the tibialis anterior muscles were dissected prior to transcardial perfusion and post-fixed in 4% PFA for 10 min after which they were rinsed in PBS and cryoprotected in 30% sucrose in PBS overnight at 4°C or until tissue has sunk. Tissues were embedded in Tissue-Tek OCT compound and frozen on dry-ice. Muscle samples were cut longitudinally at 100 μm thickness and slide mounted. For optimal tissue permeabilization, tissue was incubated with 2% Triton-X 100 in PBS for 30 minutes followed by a 30-minute block consisting of 10% normal donkey serum (NDS) and 1% Triton-X 100 in PBS. Slides were incubated with primary antibodies; presynaptic marker 2H3 (1:50, DSHB, Iowa, USA) and SV2 (1:100, DSHB, Iowa, USA) diluted in the blocking solution at room temperature. the following day, slides were washed 3 times in PBS before incubation with α-bungarotoxin conjugated to AlexaFluor 488 (1:1000, Invitrogen, Massachusetts USA) to label post-synapses, Alexa Fluor 674-conjugated cholera toxin subunit B (1:200, Invitrogen, Massachusetts USA) to label axonal gangliosides and secondary antibody AlexaFluor 555 donkey anti mouse (1:250, ThermoFisher, Massachusetts USA) in PBS for 2 hours at room temperature. Per animal, an average of 27 NMJs per TA muscle were quantified as: 1. innervated; 2. partially innervated, or 3. denervated. CTB staining was positive when >10 punctae around the NMJ along the motor terminal were observed.

### Primary cultures

Primary spinal cord lower motor neurons (MN), from SOD1^G93A^ rat embryos, which were genotyped at E14, are more sensitive to glutamatergic injury and respond less to neuroprotective compounds than the motor neurons from wild-type (WT) mice. Rat spinal cord motoneurons were prepared [38] and maintained in Neurobasal medium with a 2% solution of B27 supplement, 2nM of I-glutamine, 2% of PS solution, and 10 ng/mL of brain-derived neurotrophic factor (BDNF). On day 12 of culture, primary motor neurons were pre-treated with relevant agents, for 24 hours. On day 13 glutamate (5 μM) was added for 20 minutes, prior to washout and re-addition of agents for 24 hours. Test compounds were tested on one culture in 96 well-plates (6wells per condition). The end-points were neuronal survival (MAP2 staining), total length of neurite network (MAP-2, μm) and cytoplasmic accumulation of TDP-43, in MAP-2 +ve neurons) which accumulates in cytoplasm of motor neurons in nearly all cases of ALS. Statistical analyses were performed by one-way ANOVA, followed by Fisher’s LSD test.

### Statistics

For behavioural analysis, ANOVA was used and dependent on which test, 2 or 3-way repeated measures (RM) was used where appropriate. Interactions were followed up with a *Bonferroni post hoc* test (GraphPad Prism Software, version 9, San Diego, CA). All data were represented as mean ± SEM and *p*<0.05 was considered significant.

## Acknowledgments

This work was supported by a FightMND Translational Research Grant (B.J.T.), FightMND IMPACT Grant (S.J.L.) and Stafford Fox Medical Research Foundation Grant (B.J.T.). B.J.T was supported by a NHMRC-ARC Dementia Research Leadership Fellowship 1137024. The Florey Institute of Neuroscience and Mental Health acknowledges support from the Victorian Government, in particular, funding from the Operational Infrastructure Support Grant.

