## Supplementary Fig for "Profound lipid dysregulation in mutant TDP-43 mice is ameliorated by the glucocerebrosidase 2 inhibitor ambroxol"

**Supplementary Section**

**Supplementary methods:**

*Untargeted lipidomic analysis*

Untargeted Liquid chromatography/ mass spectrometry (LC-MS) analysis of lipids was performed as follows; The dried lipid extracts were re-suspended in 200 µl of butanol (BuOH) /MeOH (1:1) with 10 mM ammonium formate and subjected to LC-MS analysis as reported by Hu et al. (2008) [37] and described in brief below. The lipid extracts were placed in the autosampler set at 12 °C and separated by injecting 15 µl aliquots into an InfinityLab Poroshell 120 EC-C18 2.1 x 100 mm (2.7-Micron particle size) column (Agilent, USA) operated at 55 °C using an Agilent 1290 HPLC system and a flow rate of 0.26 mL/min. Elution was performed over a 30 min binary gradient consisting of acetonitrile (ACN)-water (60:40, v/v) and isopropanol (IPP)-ACN (90/10, v/v) both containing 10 mM ammonium formate as eluent A and B respectively. The gradient used was; first a 0-1.5 min isocratic elution with 32% B which was then increase to 45% B from 1.5 to 4 min, then to 52% B from 4 to 5 min followed by an increase to 58% B from 5 to 8 min. Next, it was increased to 66% B from 8 to 11 min followed by an increase to 70% B from 11 to 14 min and an increase to 75% B from 14 to 18 min. Then, from 18 to 21 min B was increased to 97% and B was maintained at 97% from 21-25 min. Finally, solvent B was decreased to 32% from 25 to 25.10 min and B was maintained at 32% for another 4.9 min for column re-equilibration [37].

Lipids were analyzed using a Sciex Triple TOFTM 6600 QqTOF mass spectrometer equipped with a Turbo VTM dual-ion source [electro-spray ionization (ESI) and atmospheric pressure chemical ionization (APCI)] and an automated calibrant delivery system (CDS) using Sequential Window Acquisition of All Theoretical Mass Spectra (SWATH-MS) in positive ion mode [38]. The parameters were set as follows; MS1 mass range: 100-1700 m/z, SWATH scan range: 300-1700 m/z, MS/MS mass range: 100-1700 m/z, time of flight (TOF) MS accumulation time: 50.0 ms, TOF MS/MS accumulation time: 10 ms, collision energy: +45 V, collision energy spread: 15 V, precursor window: 15 Da and the cycle time: 1042 ms. The following ESI parameters were used: source temperature: 250 ºC, curtain gas: 35 psi, Gas 1: 25 psi, Gas 2: 25 psi, Declustering potential: +80 V and Ion spray voltage floating: 5500 V. The instrument was calibrated automatically with the CDS delivering APCI calibration solution every five samples.

*Immunoblotting*

Lumbar spinal cord and tibialis anterior muscle were snap frozen and protein were extracted as previously described [40]. Proteins from tissue lysates (40µg) were electroporated on 12.5 % SDS polyacrylamide precast gels (Bio-Rad) after which the separated protein were transferred on Immun-Blot PVDF membrane (Bio-Rad). Membranes were blocked for 1 hr in blocking buffer consisting of 5% skim milk dried powder in Tris-buffered saline with Tween-20 (TBST; tris base 20mM, NaCl140mM, pH 8.0,0.1% Tween-20 (Sigma)). Primary antibodies against ms P62(1:500, abcam), ms/rb βactin (1:2000, Sigma), rbt VDACI (1:1000 abcam), rb LC3 (1:1000, Sigma) in 3% BSA (Sigma) in TBST were incubated O/N at 4 degrees Celsius. Membranes were washed in TBST 3 times before incubation with IRDye 680- or 800CW conjugated secondary anti mouse and rabbit antibodies (1:10000, Li-Cor biosciences) in TBST. Membranes were visualised using the chemidoc imaging system (Bio-Rad). Band intensity was normalised to internal βactin and wildtype vehicle was used as control. Band intensity was quantified using ImageJ (National Institutes of Health, USA, version 1.15s).

*SCoRE imaging*

Spectral Confocal Reflectance Microscopy was employed on spinal cord sections as described by [41]. The intensity of myelin using SCoRE in the corticospinal tract of longitudinal lumbar spinal cord sections was assessed at P60 and P210, corresponding with our lipidomic analysis. Regions of interest (ROIs) were 100x100 pixels and intensity was quantified using ImageJ (National Institutes of Health, USA, version 1.15s). Per animal, 5 ROIs were assessed.

### Supplementary Figures and Tables

**Supplementary Figure 1. Raw peak data analysis of main lipid species in TDP-43$^{Q331K}$ mice**. Analysis of raw peak area data of lipids in spinal cord (A) shows significant higher abundance of several subtypes of lysophosphatidylcholine (LPC), Phosphatidylcholine (oxidised and non-oxidised) (PC), Phosphatidylethanolamine (PE), Phosphatidylglycerol (PG), Phosphatidylinositol (PI) and triglycerides (TG). In skeletal muscle (B), similar subtypes, apart from TG, are observed to be more abundant in the TDP-43$^{Q331K}$ mice, *n*=6/group.

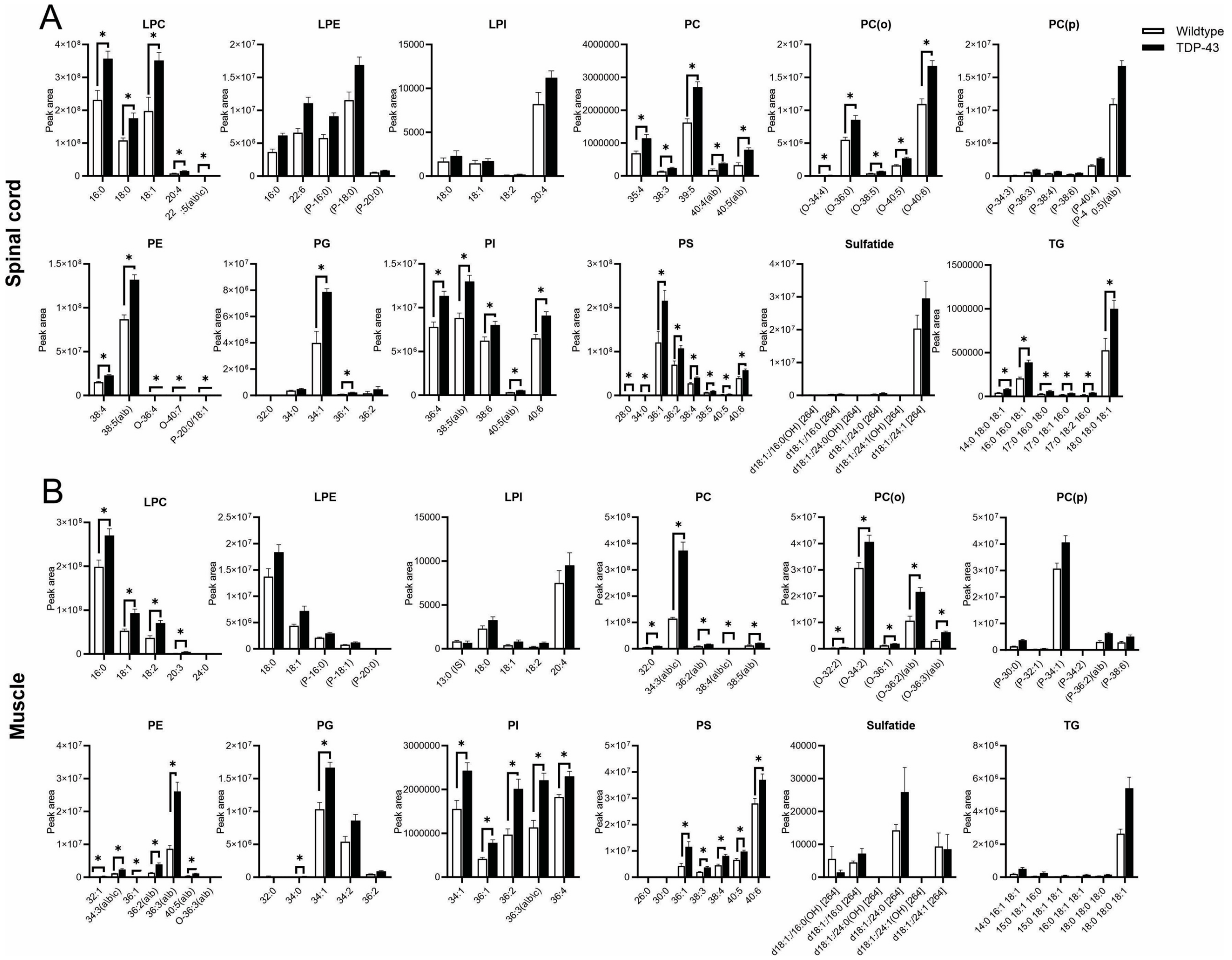

**Supplementary Figure 2. Untargeted lipidomic analysis of TDP-43$^{Q331K}$ mouse tissue.** Principal component analysis of untargeted lipidomics in lumbar spinal cord (A) and skeletal muscle (D) shows larger variability in WT spinal cord, where WT muscle lipidome is relatively tightly clustered. Volcano plot based on t-Test with fold change threshold (X-axis) of 1.5 and *p*-value cut-off at 0.05 (Y-axis) coloured circles represent lipids above these thresholds, in the spinal cord there were 17 decreased and 63 increased lipids in Wildtype vs. TDP (B), where in the Tibialis Anterior muscle 113 were decreased and 97 increased (F). Heatmap of top 50 lipids identified through untargeted LCMS in spinal cord (C) and TA muscle (F) shows clear clustering based on genotype suggesting clear lipid restructuring in the TDP-43 spinal cord *and* skeletal muscle, *n*=6/group.

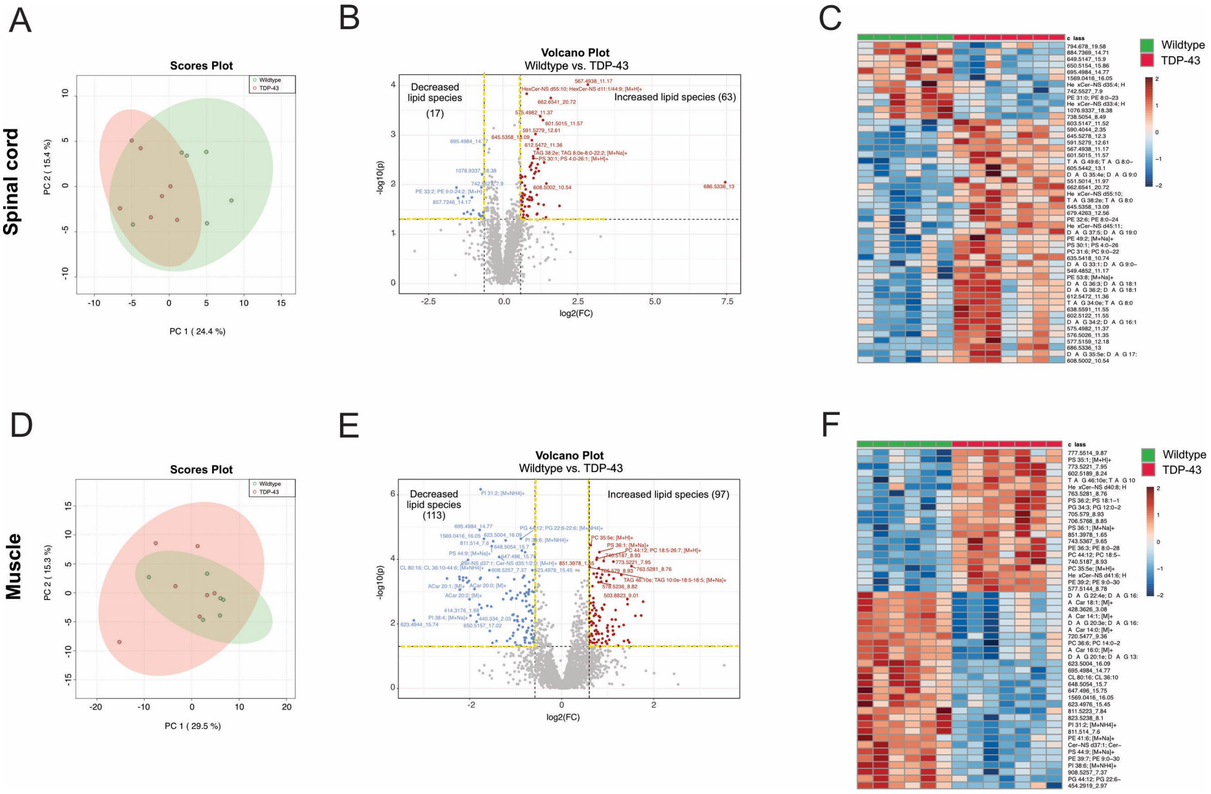

**Supplementary Figure 3. Autophagy and myelin density is unaffected in TDP-43$^{Q331K}$ mice.** Representative blot of protein analysis in spinal cord at P300 (A), shows no differences in autophagy marker P62 and VDAC1 (B) or LC3II/LC3I ratio (C) between wildtype and TDP-43 mice. Further, no effect of Ambroxol on autophagy was observed. Gene expression of autophagy markers in the spinal cord also showed no significant effects (D). Gene expression of denervation markers MUSK and Atrogin seem to be unaffected in the gastrocnemius muscle at P300 (E). Intensity of Spectral Confocal Reflectance Microscopy (SCoRe) (F) of the corticospinal tract (CST) longitudinally at P60 and P210 showed no significant difference in myelin density between genotype or Ambroxol treatment (G). Data represented as mean ± SEM, *n*=5-8/group.

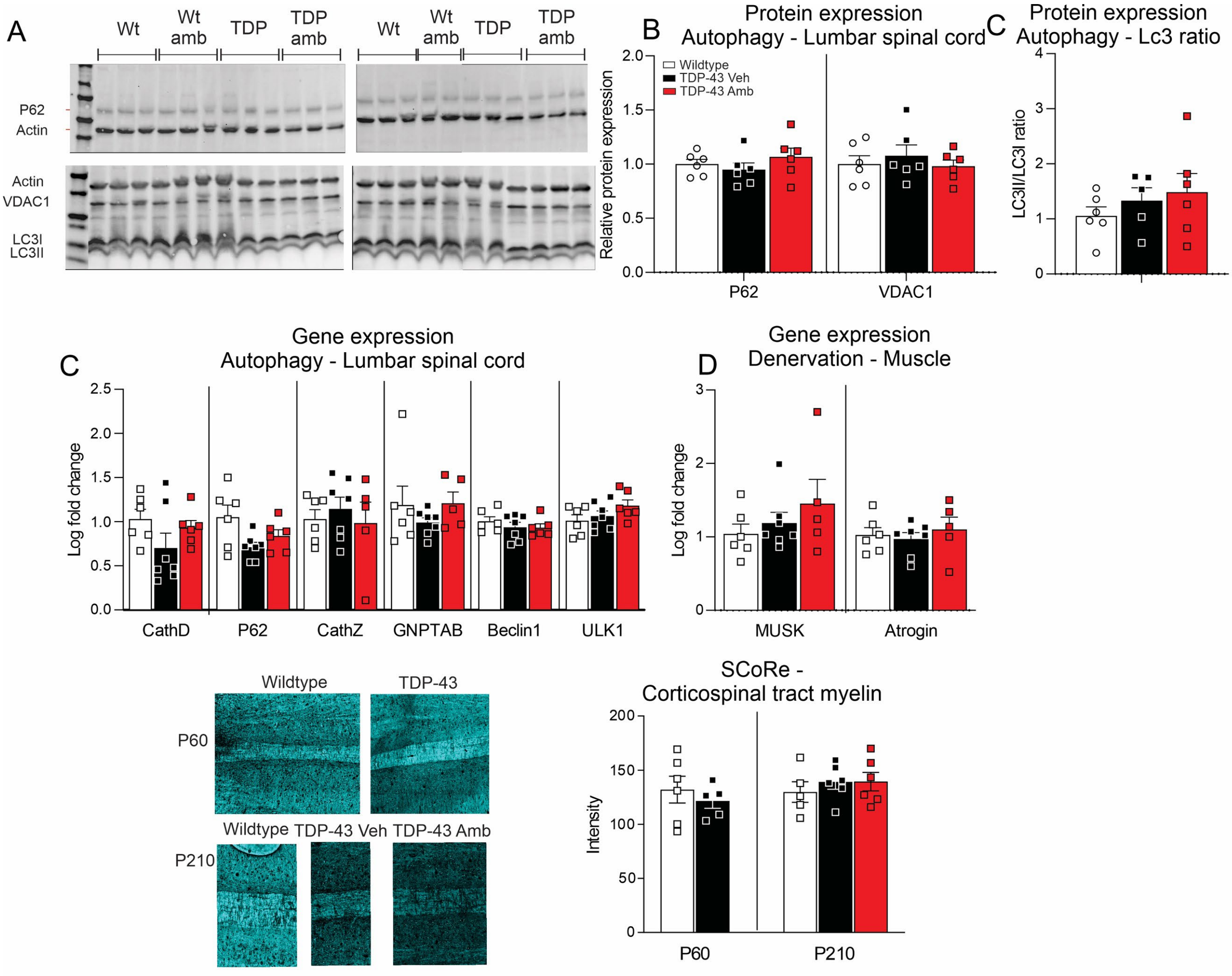

**Supplementary Figure 4. Untargeted lipidomic analysis of the effect of ambroxol on spinal cord and tibialis anterior muscle of TDP-43$^{Q331K}$ mice.** Principal component analysis of untargeted lipidomics in lumbar spinal cord (A) and skeletal muscle (D) shows large variability in Ambroxol treated tissue, where untreated tissue is more tightly clustered. Volcano plot based on t-Test with fold change threshold (X-axis) of 1.5 and *p*-value cut-off at 0.05 (Y-axis) coloured circles represent lipids above these thresholds, in the spinal cord there were 5 decreased and 25 increased lipids in vehicle vs Ambroxol treated spinal cord tissue (B), where in the Tibialis Anterior muscle 7 were decreased and 33 increased (F). Heatmap of top 50 lipids identified through untargeted LCMS in spinal cord (C) and TA muscle (F) shows clear clustering based on treatment suggesting Ambroxol affects the lipidome in both CNS and skeletal muscle of TDP-43$^{Q331K}$ mice, *n*=6/group.

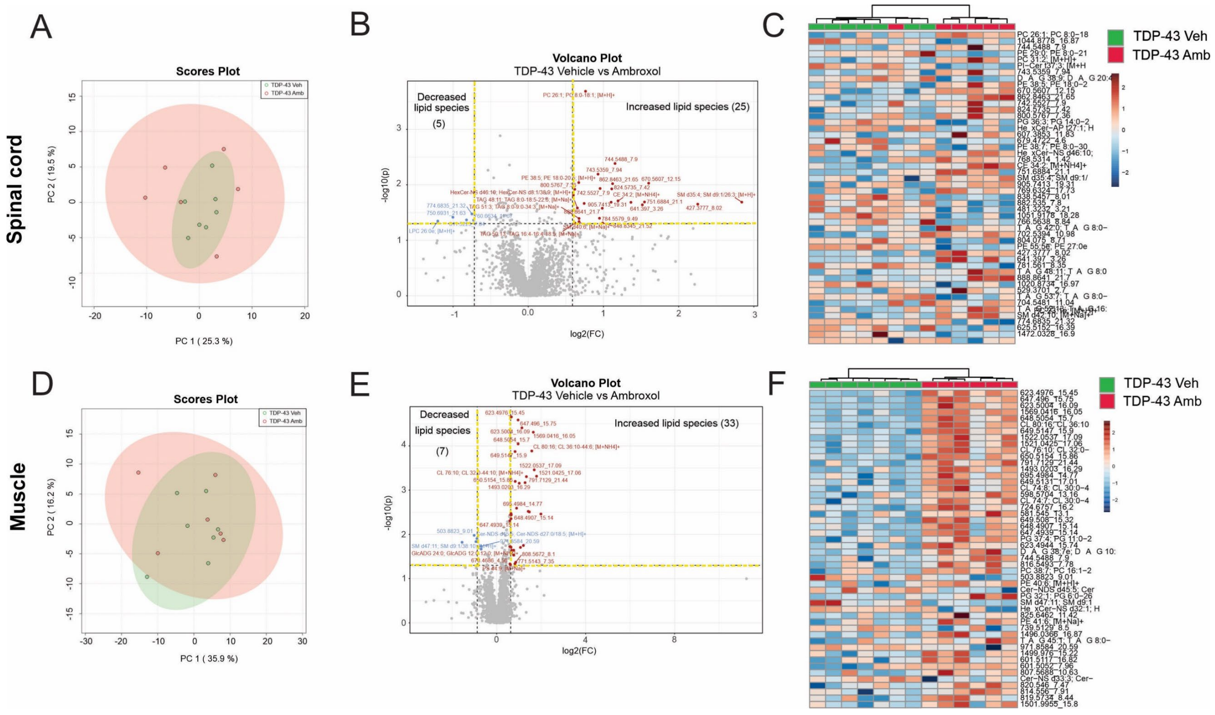

**Supplementary Table 1** Gait analysis of the fore limbs in TDP-43 and Wildtype mice on the digigait. *indicates $p<0.05$.

| Fore limbs - Digigait statistics - ANOVA main genotype effect | | | | | | | | |
|---|---|---|---|---|---|---|---|---|
| **Variable** | P60 | P90 | P120 | P150 | P180 | P210 | P240 | P270 |
| Swing | | * | * | * | | * | | * |
| %SwingStride | | | | | | | | |
| Brake | | | | | | | | |
| %BrakeStride | | | | | | | | |
| Propel | | | * | | | | * | * |
| %PropelStride | | | | | | | | |
| Stance | | * | * | * | | | * | * |
| %StanceStride | | | | | | | | |
| Stride | | | * | * | | | * | * |
| %BrakeStance | | | | | | | | |
| %PropelStance | | | | | | | | |
| Stance/Swing | | | | | | | | |
| StrideLength | | * | * | * | | | * | * |
| Stride Frequency | | * | * | * | | | * | * |
| PawAngle | | | | | | | | * |
| Absolute PawAngle | | | | | | | | |
| Paw Angle Variability | | | * | | * | | | * |
| StanceWidth | * | | * | | | | | |
| StepAngle | | | | | | | | * |
| SLVar | | | | | | | | |
| SWVar | | | | | | * | | |
| StepAngleVar | | | | | | | | |
| #Steps | | | | | | | | * |
| Stride Length CV | | | * | | * | * | * | |
| Stance Width CV | * | | * | | | | | |
| Step Angle CV | | | | | | | | |
| Swing Duration CV | | | * | | | | | |
| Paw Area at Peak Stance in sq. cm | * | | | * | | | * | |
| Paw Area Variability at Peak Stance in sq. cm | | | * | | | | | |
| Hind Limb Shared Stance Time | | | | | | | | |
| % Shared Stance | | | | | | | | |
| StanceFactor | | | * | | | | | * |
| Gait Symmetry | | | | | | | | |
| MAX dA/dT | | | | | | | | |
| MIN dA/dT | * | | | | | | | |
| Tau - Propulsion | | | | | | | | |
| Overlap Distance | * | * | * | * | * | * | * | |
| PawPlacementPositioning[PPP] | | | * | | | | * | |
| Ataxia Coefficient | | | * | | | * | * | * |
| Midline Distance | | * | * | * | * | * | * | * |
| Axis Distance | | | | | | | | |
| Paw Drag | | | | | | | | |

**Supplementary Table 2** Gait analysis of the fore limbs in TDP-43 treated with and without ambroxol on the digigait. *indicates $p<0.05$.

| Fore limbs - Digigait statistics - ANOVA genotype X drug effect | | | | | | | | |
|---|---|---|---|---|---|---|---|---|
| **variable** | P60 | P90 | P120 | P150 | P180 | P210 | P240 | P270 |
| Swing | | * | | | | * | * | |
| %SwingStride | | | | | | * | | |
| Brake | | | | | | | | |
| %BrakeStride | | | | | | | | |
| Propel | | | | | | | | |
| %PropelStride | | | | | | * | | |
| Stance | | | | | | | | |
| %StanceStride | | | | | | * | | |
| Stride | | | * | | | | | |
| %BrakeStance | | | | | | | | |
| %PropelStance | | | | | | | | |
| Stance/Swing | | | | | | * | | |
| StrideLength | | | * | | | | * | |
| Stride Frequency | | | * | | | | * | |
| PawAngle | | | | | | | | |
| Absolute PawAngle | | | | | | | | |
| Paw Angle Variability | | | | | | | | |
| StanceWidth | | | | | | | | |
| StepAngle | | | | | | | | |
| SLVar | | | | | | | | |
| SWVar | | | | * | | | | |
| StepAngleVar | | | | * | | | | |
| #Steps | | * | | * | | | | * |
| Stride Length CV | | | | | | * | | |
| Stance Width CV | | | | | | | | |
| Step Angle CV | | | | * | | | | |
| Swing Duration CV | | * | | | | | | |
| Paw Area at Peak Stance in sq. cm | | | | | | | * | |
| Paw Area Variability at Peak Stance in sq. cm | | | | | | | | |
| Hind Limb Shared Stance Time | | | | | | | | |
| % Shared Stance | | | | | | | | |
| StanceFactor | | * | | | | | | |
| Gait Symmetry | | | | | | | | |
| MAX dA/dT | | | | | | | | |
| MIN dA/dT | | * | | | | | | |
| Tau - Propulsion | | | | | | | | |
| Overlap Distance | | | | | | | * | * |
| PawPlacementPositioning[PPP] | | | | | | | | |
| Ataxia Coefficient | | | | | | | | |
| Midline Distance | | | | | | * | | |
| Axis Distance | | * | | | | | * | |
| Paw Drag | | | | | | | | |

**Supplementary Table 3** Gait analysis of the hind limbs in TDP-43 and Wildtype mice on the digigait. *indicates $p<0.05$.

| **Hind limbs - Digigait statistics - ANOVA main genotype effect** | | | | | | | | |
|---|---|---|---|---|---|---|---|---|
| **variable** | P60 | P90 | P120 | P150 | P180 | P210 | P240 | P270 |
| Swing | | | * | | | | | * |
| %SwingStride | | | | | | | | |
| Brake | | | | | | | | |
| %BrakeStride | | | * | | | | | |
| Propel | | | * | * | | * | * | * |
| %PropelStride | | | | | | * | | |
| Stance | | * | * | * | | | * | * |
| %StanceStride | | | | | | | | |
| Stride | | * | * | * | | | * | * |
| %BrakeStance | | | * | | | | | |
| %PropelStance | | | * | | | | | |
| Stance/Swing | | * | | | | | * | |
| StrideLength | | * | * | * | | | * | * |
| Stride Frequency | | * | * | * | | | * | * |
| PawAngle | | | * | * | | | | * |
| Absolute PawAngle | | | | * | | | | |
| Paw Angle Variability | | | * | | | | | |
| StanceWidth | | | | | * | * | * | |
| StepAngle | | | | | | | | |
| SLVar | | | * | | | | | * |
| SWVar | | | | | | | | |
| StepAngleVar | | | | | | | | |
| #Steps | | | | | | | | |
| Stride Length CV | | | * | | | | * | * |
| Stance Width CV | | | | | | | | |
| Step Angle CV | | | | | | | | |
| Swing Duration CV | | | * | | | | | |
| Paw Area at Peak Stance in sq. cm | | | | | | * | | |
| Paw Area Variability at Peak Stance in sq. cm | | | | | | | | |
| Hind Limb Shared Stance Time | * | * | | * | | | * | * |
| % Shared Stance | | | | * | | | | |
| StanceFactor | | * | | * | | | | |
| Gait Symmetry | | | | | | | | |
| MAX dA/dT | | | | | | * | | |
| MIN dA/dT | * | | | * | | * | * | |
| Tau - Propulsion | | | | | | | * | * |
| Overlap Distance | * | * | * | * | * | * | * | |
| PawPlacementPositioning[PPP] | | | * | * | | | * | |
| Ataxia Coefficient | | | * | | | | | * |
| Midline Distance | | | | * | | | * | |
| Axis Distance | | | | | | | | |
| Paw Drag | * | * | * | * | * | * | * | * |

**Supplementary Table 4** Gait analysis of the hind limbs in TDP-43 treated with and without ambroxol on the digigait. *indicates  p<0.05.

| **Hind limbs - Digigait statistics - ANOVA genotype X drug effect** | | | | | | | | |
|---|---|---|---|---|---|---|---|---|
| **variable** | P60 | P90 | P120 | P150 | P180 | P210 | P240 | P270 |
| Swing | | * | * | | | * | * | |
| %SwingStride | | | | | | | | |
| Brake | | | | | | | | |
| %BrakeStride | | | | * | | | | |
| Propel | | | | | | | | |
| %PropelStride | | | | | | | | |
| Stance | | | | | | | | |
| %StanceStride | | | | | | | | |
| Stride | | | | | | * | * | |
| %BrakeStance | | | | | | | | |
| %PropelStance | | | | * | | | | |
| Stance/Swing | | | | | | | | |
| StrideLength | | | * | | | | * | * |
| Stride Frequency | | | | | | | * | |
| PawAngle | | | | | | | | |
| Absolute PawAngle | | | | | | | | |
| Paw Angle Variability | | | | | | | | |
| StanceWidth | | | | | * | | | |
| StepAngle | | | | | | | * | |
| SLVar | | | | | | | | |
| SWVar | | | | | | | | |
| StepAngleVar | | | | | | | | |
| #Steps | | * | | | | * | | |
| Stride Length CV | | | | | | | * | |
| Stance Width CV | | | | | | | | |
| Step Angle CV | | | | | | | * | |
| Swing Duration CV | | | | | | | | |
| Paw Area at Peak Stance in sq. cm | | | | | * | | | |
| Paw Area Variability at Peak Stance in sq. cm | | | | | | | | |
| Hind Limb Shared Stance Time | | | | | | | | |
| % Shared Stance | | | | | | | | |
| StanceFactor | | * | | | | | * | |
| Gait Symmetry | | | | | | | | |
| MAX dA/dT | | | | | | | | |
| MIN dA/dT | | | | | * | * | | |
| Tau - Propulsion | | | | | | * | | |
| Overlap Distance | | | | | * | | * | * |
| PawPlacementPositioning[PPP] | | | | | | | | |
| Ataxia Coefficient | | | | | | | * | * |
| Midline Distance | | | | | | * | | |
| Axis Distance | | | | | | | | |
| Paw Drag | | | | | | | | |

**Supplementary Table 5**. main lipid species and the amount of subspecies identified in targeted lipidomic analysis of wildtype and TDP-43$^{Q331k}$ tissue, their function and literature reference observed in previous literature in relevance to ALS.

| Lipid name and abbreviation | # subspecies | Main function | Up or down in ALS (ref) |
|---|---|---|---|
| AcylCarnitine (Acar) | 13 | Fatty acid transport to mitochondria | ↓ [44] |
| Cholesteryl ester (CE) | 26 | Lipid storage and metabolism | ↑[27] |
| Ceramide (Cer) | 42 | Core structure of sphingolipids | ↑[27] |
| Unesterified Cholesterol (COH) | 2 | Cholesterol synthesis and in turn cell membrane health | unknown |
| Diacylglycerol (DG) | 21 | Lipid biosynthesis, protein kinase C activator (PKC) | ↑PKC [45] |
| Dihydroceramide (DhCer) | 7 | Precursor of ceramide | ↑ [7] |
| Ganglioside GM1 (GM1) | 1 | Nerve conduction and synaptic transmission | ↑[7, 40] |
| Ganglioside GM3 | 7 | Neuronal proliferation | ↑[40] |
| Hexa-ceramide (Hex1Cer, Hex2Cer, Hex3Cer) | 33 | Possible metabolic disease marker | unknown |
| Lyso-phosphatidylcholine (LPC) | 38 | Increased in ischemic injury (koizumi et al 2010) | Unknown |
| Lyso-phosphatidylethanolamine (LPE) | 12 | Membrane phospholipid function | Unknown |
| Lyso- phosphatidylinositol (LPI) | 5 | Neuronal survival signalling | ↑PI3-Kinase activity [46, 47] |
| Oxidised cholesteryl ester (oxCE +2O NH4) | 2 | Cholesterol trafficking through low density lipoprotein (LDL) | Unknown |
| Phosphatidylcholine (PC) | 72 | Acetylcholine synthesis | ↑[48] |

| | | | |
|---|---|---|---|
| Phosphatidylethanolamine (PE) | 68 | Membrane structure and function | ↑[49] |
| Phosphatidylglycerol (PG) | 10 | Membrane structure and function | Unknown |
| Phosphatidylinositol (PI) | 16 | Membrane trafficking and lipid signalling | ↑PI3-Kinase activity [46, 47] |
| Phosphatidylserine (PS) | 13 | Healthy nerve cell membranes and myelin/mitochondrial function | ↑[7] |
| Sphingomyelin (SM) | 35 | Membrane homeostasis | ↑[48] |
| Sulfatide | 7 | Major glycolipid component of myelin | ↑[50] |
| Triacylglycerol (TG) | 48 | Lipoprotein metabolism | ↓[48] |
